# A Multivalent Pan-Ebolavirus Nanoparticle Vaccine Provides Protection in Rodents from Lethal Infection by Adapted Zaire and Sudan Viruses

**DOI:** 10.1101/2025.01.29.635581

**Authors:** Natalie Brunette, Connor Weidle, Samuel P Wrenn, Brooke Fiala, Rashmi Ravichandran, Kenneth D Carr, Samantha E Zak, Elizabeth E Zumbrun, Russell R Bakken, Michael Murphy, Sidney Chan, Rebecca Skotheim, Andrew J Borst, Lauren Carter, Colin E Correnti, John M Dye, David Baker, Neil P King, Lance J Stewart

**Affiliations:** Department of Biochemistry, University of Washington, Seattle, WA 98195, USA; Institute for Protein Design, University of Washington, Seattle, WA 98195, USA; Archon Biosciences, Seattle, WA 98102, USA; Bonum Therapeutics, Seattle, WA 98102, USA; AstraZeneca, Icosavax Inc., Seattle, WA 98102, USA; US Army Medical Research Institute for Infectious Disease (USAMRIID), Fort Detrick, MD, USA; Pfizer, Seagen, Seattle, WA 98102, USA; Gates MRI, Seattle, WA 98102, USA; Clinical Research Division, Fred Hutchinson Cancer Research Center Seattle, WA 98109, USA; Howard Hughes Medical Institute, University of Washington, Seattle, WA, USA

## Abstract

Both *Zaire ebolavirus* (EBOV) and *Sudan ebolavirus* (SUDV) are members of the family *Filoviridae*, first discovered in 1976 during outbreaks of hemorrhagic fever in northern Zaire and southern Sudan. Ebola virus disease outbreaks are major public health events because of their potential for human-to-human transmission with high case fatality rates. Filoviral surface glycoproteins (GPs) are known to be the primary targets of neutralizing antibodies for protection from disease, and are the relevant immunogens in the two approved EBOV vaccines. Here we describe the design, electron microscopy-based structural characterization, and efficacy testing of a series of icosahedral I53-50 nanoparticles displaying prefusion trimeric EBOV and SUDV GP antigens. Mice and guinea pigs vaccinated with either a cocktail of EBOV-GP-I53-50 plus SUDV-GP-I53-50 or mosaic EBOV / SUDV-GP-I53-50 nanoparticles were protected from death or severe clinical signs of disease and weight loss, respectively, when challenged with either mouse-adapted EBOV or guinea pig-adapted SUDV.

## Introduction

The *Ebolavirus* genus is a member of the single-stranded negative-sense RNA virus family *Filoviridae* and includes both *Zaire ebolavirus* (EBOV) and *Sudan ebolavirus* (SUDV). The study of filoviruses and the development of vaccines against this family has been difficult, in part due to the high biosafety level required for research. Ebolaviruses were first discovered after 1976 outbreaks in southern Sudan (now South Sudan) and Zaire (now the Democratic Republic of Congo), and have remained a major public health issue. The most common Ebola virus disease (EVD^1^) outbreaks are caused by EBOV, with over 20 outbreaks in Sub-Saharan Africa having case-fatality rates ranging from 40%-70%^2,3^. The largest ever outbreak of Ebola occurred in West Africa from 2013–2016, which resulted in over 28,000 cases and over 11,000 deaths. In 2022 SUDV also re-emerged, causing more than 160 confirmed or probable cases^4^ with 55 confirmed deaths^5^. Considering the life-threatening and disruptive nature of EVD outbreaks, there is a tremendous need for effective vaccines that provide broad-spectrum prophylactic protection against both EBOV and SUDV.

Several EBOV and SUDV vaccine candidates have been shown to protect against lethal infection in non-human primates (NHPs)^6–8^. Currently, the two licensed Ebola vaccines are approved only for prevention of EVD caused by EBOV^9^. Merck’s Ervebo® was the first and remains the only EVD vaccine to be approved by the Food and Drug Administration (FDA) in the United States (November 2019). It was developed by the Public Health Agency of Canada, and later licensed to Merck for further development and manufacturing. The vaccine, also known as rVSV-ZEBOV, is a replication-competent, live-attenuated recombinant vesicular stomatitis virus (VSV) vaccine that expresses the main trimeric GP from *Zaire ebolavirus*^8,10^. In 2015, over two thousand close contacts of confirmed Ebola patients were ring-vaccinated with rVSV-ZEBOV on an emergency basis during the outbreak in West Africa to reduce the spread of the infection^11^. Following a large-scale ring vaccination effort in 2019, the WHO published preliminary results stating that the vaccine was 97.5% effective at preventing transmission of the virus^12^, including in one case of a healthcare worker presenting with a late reactivation of EVD in the United Kingdom. Additionally, voluntary contact-based vaccination with rVSV-ZEBOV demonstrated that none of the 25 people exposed to the virus and vaccinated with rVSV-ZEBOV experienced EVD^13^.

Janssen Pharmaceutica together with Bavarian Nordic developed the two-dose Zabdeno/Mvabea (Ad26.ZEBOV/MVA-BN-Filo) vaccine that was the second vaccine to be licensed (July 2020); at this time only by the European Medicines Agency (EMA) for prevention of EVD caused by *Zaire ebolavirus*. Zabdeno/Mvabea uses a prime with Ad26.ZEBOV (Zabdeno) followed 8 weeks later by a heterologous boost with MVA-BN-Filo boost (Mvabea)^9^. Zabdeno is a human adenovirus serotype 26 encoding Zaire Mayinga EBOV-GP. Mvabea is a modified Vaccinia Ankara virus (MVA) encoding EBOV GP, SUDV GP, Marburg virus (MARV) GP, and Taï Forest virus (TAFV) nucleoprotein. Noted disadvantages of this approach include the possibility of preexisting immunity to the Ad26 or MVA vectors, which may reduce the effectiveness of the vaccine, as well as the 8 week wait period before the second dose is administered which is not ideal for outbreak settings.

That the two approved EVD vaccines are both designed to produce GP immunogen(s) provides substantial evidence that filoviral GPs as immunogens can serve as prophylactic vaccines for EVD. However, There is currently no clinically validated single vector or subunit vaccine that offers broad protection against more than one strain of EVD-causing viruses. Numerous studies have shown that obtaining cross-protection against both EBOV and SUDV with a single GP based vaccine is challenging. In humans infected with either EBOV or SUDV, or vaccinated against EBOV with 1 or 2 doses of the rVSV-ZEBOV vaccine, antibody cross-reactivity between stabilized EBOV and SUDV GP trimers^14^ occurs only occasionally and typically with markedly reduced binding and neutralization titres^15^. In repurposed NHPs from a successful rVSV-EBOV vaccine efficacy study with EBOV challenge, there was little cross-protection to SUDV challenge even though animals were found to develop some cross-reactive humoral responses to SUDV^6,16^. In contrast, a single-dose vaccination with SUDV GP vaccine (rVSV-SUDV) protected NHPs from lethal infection by the SUDV strain SUDV-Gulu^16^. The lack of heterologous cross-viral EVD protection from GP based vaccines is likely due to the significant amino acid sequence differences between EBOV and SUDV GPs which have 55% amino acid sequence identity over their full open reading frames (**Supplemental Figure 1**).

Nanoparticle (NP) or virus-like particle (VLP) platforms have gained traction in modern vaccine design, as presenting antigen in a repetitive array improves antigen presenting cell uptake and B-cell activation through avidity effects^17,18^. The first licensed computationally designed protein medicine is SKYCovione™, a self-assembling NP vaccine invented in 2020^19–21^ during the COVID-19 pandemic. SKYCovione™ is composed of a robust self-assembling two-component icosahedral chassis called the I53-50 NP system^22^. It is constructed from 20 trimeric (“I53-50A”) and 12 pentameric (“I53-50B”) subunits that can be separately produced and then mixed in solution to cooperatively self-assemble into a uniform 120-subunit icosahedral nanoparticle^23^. The SKYCovione™ vaccine for SARS-CoV-2 (approved in S. Korea and the UK)^20,21^ displays 60 copies of the SARS-CoV-2 spike glycoprotein (S) receptor binding domain (RBD) as a genetic fusion to the trimeric I53-50A component^19^. Inspired by pioneering work in the co-display of multiple influenza viral hemagglutinin (HA) RBD antigens in a mosaic nanoparticle format^24^, studies using the I53-50 NP to co-display multiple sarbecovirus RBDs as a mosaic or cocktail demonstrated protection against SARS-CoV challenge in mice even in the absence of the SARS-CoV RBD in the vaccine^25^. These studies and others^26,27^ establish a growing body of evidence that by deploying computational protein design, it is possible to provide the immune system with multivalently displayed immunogens in a nanoparticle format to elicit desired immune responses for broad spectrum antiviral prophylaxis. While protein nanoparticle-based Ebola vaccine candidates have been reported using full-length GP purified from insect cells^28,29^, ferritin nanoparticles^30^ or the computationally designed I3-01 nanoparticle chassis^31,32^, these nanoparticle systems lack the ability to easily co-display multiple antigens in a spatially controlled manner, which can be readily accommodated by the two-component I53-50 NP chassis.

Since multivalent mosaic immunogen-NPs can induce heterotypic prophylaxis^24–27^, they offer the potential to achieve broad-spectrum (viral family) protection from vaccination. Such mosaic NP immunogens can also be used to elicit and discover broadly neutralizing antibodies (bNAbs)^25,33^. Due to its cyclic C3 symmetry, the trimeric subunit I53-50A is ideally suited to genetic fusion to the C terminus of homo-trimeric viral glycoproteins^25,26,34^. Moreover, the underlying I53-50 NP chassis can lead to the stabilization of displayed trimeric glycoprotein immunogens^35^. The improved solution behavior of trimeric glycoprotein fusions to the I53-50A component affords a facile manufacturing paradigm where ratios of different immunogens can in principle be rapidly formulated with the I53-50B pentameric subunit into a single qualified NP vaccine product for broad-spectrum vaccine formulation. Here, we leverage the multi-component nature of the I53-50 NP chassis to develop mosaic and cocktail nanoparticle vaccine candidates displaying EBOV (Zaire Mayinga) and SUDV (Sudan Gulu) GPs. To ensure the integrity and proper assembly of each GP immunogen, we implemented a structured electron microscopy-based characterization workflow, confirming that the antigens were properly folded before and after genetic fusion to nanoparticle components, as well as following assembly into fully formed nanoparticles. Using a prime-boost vaccination strategy, we demonstrate that a cocktail of EBOV-GP-I53-50 plus SUDV-GP-I53-50; or mosaic EBOV / SUDV-GP-I53-50 immunogens formulated with AddaVax™ adjuvant elicit robust immune responses. These responses confer protection in rodent models against severe EVD following challenge with mouse-adapted EBOV (maEBOV) and guinea pig-adapted SUDV (gpSUDV), highlighting their potential as broad-spectrum vaccine candidates against this deadly class of viruses.

## Results

### Prefusion EBOV Glycoprotein Immunogen Design, Production and Characterization

Based on previous experience in our labs on the design of prefusion viral glycoprotein trimer immunogens as genetic fusions to the trimeric component of the I53-50 NP system, we elected to take a step-wise approach to the design and characterization of EBOV and SUDV GP immunogens. We first considered the optimal design of the EBOV and SUDV GP antigens on their own, prior to fusing them to the I53-50A trimer component of the I53-50 NP chassis. In this endeavor, we carefully considered the GP sub-domains that should or should not be incorporated into GP constructs. Notably, the post-Golgi full-length Ebolaviral GP (∼160 kDa) contains both N- and O-linked carbohydrate modifications and is further processed by furin proteolysis at the plasma membrane^36^, finally presenting itself on the surface of enveloped virions as trimers of heterodimers GP1 (∼140 kDa) and GP2 (∼27 kDa) (**Figure 1a**). The GP1 and GP2 units are linked together through a disulfide bond between cysteine residues C53 on GP1, which contains the receptor binding region (RBR), and C603 on GP2^37^ which contains the protein machinery responsible for the fusion of the viral and host cell membranes^38^.

**Fig. 1:**
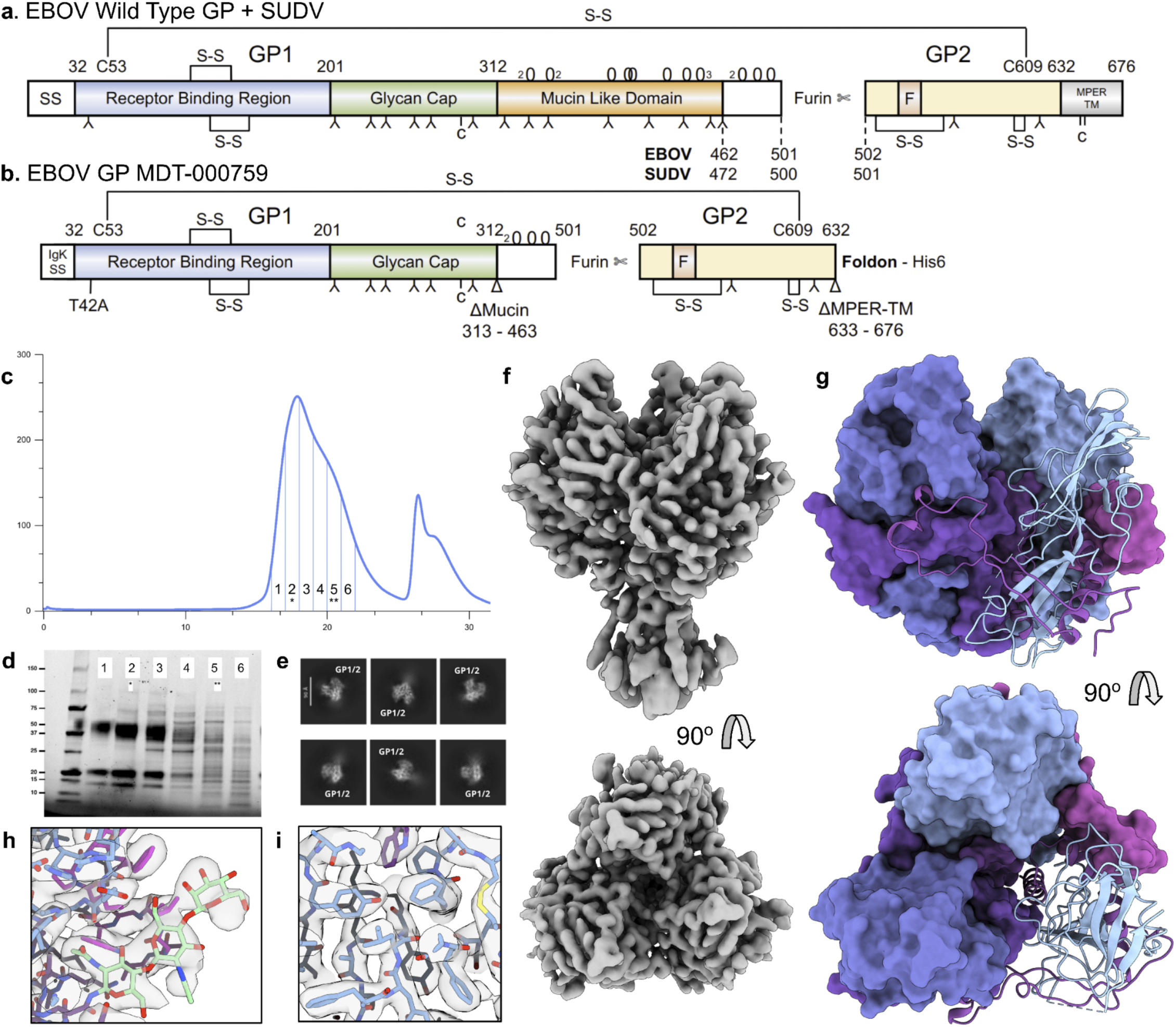
Domain Architecture of Mature Ebolavirus Glycoprotein (GP) and CryoEM Structural Characterization of EBOV-GP-Foldon. **a.** The primary wild type protein architecture for mature EBOV and SUDV GPs are illustrated with amino acid numbers for main domain boundaries noted. Glycosylation only shown for EBOV^40^ **b.** Using the same nomenclature as in **a.**, the EBOV-GP-Foldon construct (MDT-000759) domain architecture is illustrated with its deletion of the mucin-like (ΔMuc) and the MPER-TM (ΔMPER-TM) domains with C-terminal genetic fusion to the foldon domain followed by six histidine residues (His6) to facilitate IMAC purification. This construct also has the T42A mutation to knock out the first N-linked glycosylation similar to PDB 5JQ3 for EBOV (Zhao et al. 2016). Nomenclature for **a. and b.** is as follows; secretion signal peptide (SS) or IgK SS which is removed during translocation through the endoplasmic reticulum, the receptor binding region (blue), a glycan cap (green), and the mucin-like domain (orange) in GP1 which is separated from GP2 (yellow) by a furin cleavage site with fusion (F) domain (pink), membrane proximal external region (MPER) followed by the transmembrane (TM) (grey) and covalent modifications of disulfide bonds (S-S), N-linked (upside down Y), O-linked (O) (2,3 notation denote multiple neighboring O glycan sites), or C-linked glycosylation ^40,91^. **c.** EBOV-GP-Foldon was produced as secreted protein from the pCMV/R vector by transient transfection of ExpiHEK293F cells and purified by IMAC and then SEC (see Methods and **Supplemental Figure 5**) **d.** SDS polyacrylamide gel electrophoresis (SDS-PAGE), where fractions 1 and 2 were combined and used for CryoEM studies. **e.** CryoEM 2D class averages for EBOV-GP-Foldon. **f.** The 3.05 Å CryoEM map, non uniform refinement map CryoSPARC, DeepEMhancer sharpened. **g.,** Refined CryoEM structure. **h.** Build structure fit to density on the internal region. **i.** Build structure fit to density on the external region only, showing a resolved glycan at N563.

Since the heavily glycosylated mucin-like domain (muc) is thought to act as a “glycan shield” or decoy epitope for non-neutralizing antibodies^39,40^, and may even serve as an epitope for antibody-dependent enhancement of infection^41^, we first sought to design “Δmuc” deleted versions of the GPs that lacked the mucin-like domain. Furthermore, an antibody response against the mucin domain may be non-neutralizing for viral infection since the mucin domain is proteolytically removed from the virion surface during the infection process^42,43^. In addition, to ensure secretion of the constructs from mammalian cells, we removed the membrane proximal external region (MPER) and transmembrane (TM) helix^44^ from the C terminus of GP2 (ΔMPER-TM). Finally, rather than stabilize prefusion GPs with designed point mutations ^14^ or compromise prefusion GP stability using the Mano River 2014 strain sequence^45^ (**Supplemental Figure 2**), we sought to preserve as much as possible the native ancestral (Zaire Mayinga or Sudan Gulu) sequences of our Δmuc, ΔMPER-TM GP constructs. We stabilized them in the trimeric, prefusion conformation through C-terminal fusion to the “foldon” domain of T4 fibritin, which is a commonly used strategy for stabilization of class I viral fusion proteins^46,47^. Finally, guided by available X-ray crystallographic studies^47–49^, we included a T42A mutation to eliminate an N-linked glycosylation site at Asn40^40^, which has been shown to increase sample homogeneity^47,48^. This EBOV-GP-Foldon is also named MDT-000759 (**Figure 1b**).

We explored two different expression systems for the EBOV-GP-Foldon (MDT-000759): small-scale production in HEK293 cells by lentiviral transduction^50^ (**Supplementary Figure 3**), and transient transfection of ExpiHEK293F cells with pCMV/R vectors (**Figure 1c,d**). Examining the protein produced through transient transfection by cryoEM showed that the protein was a well-folded homo-trimer (**Figure 1e-i**). The structure of EBOV-GP-Foldon was determined at 3.05 Å resolution at physiological pH 7.5 (PDB ID: 9MHA, EMDB ID: EMD-48271, **Supplementary Figures 4 and 5, and Table S1**). To our knowledge, this structure provides the first high-resolution cryoEM structure of an EBOV-GP-Foldon without additional proteins bound. Overall our construct was well folded and the globular core of GP resolved well. Our EBOV-GP-Foldon structure aligned well with a high-resolution crystal structure that shares the same amino acid sequence (PDB ID: 5JQ3; all-atom RMSD of 0.918).

The glycan cap (residues 224-310) of EBOV-GP-Foldon was unresolved in our cryoEM structure. It is known that there is a cathepsin cleavage site at Lys190, which removes the glycan cap and mucin domains to allow viral entry. However, there was clear density for residues 191-195 and 216-223, suggesting that there is no off-target cleavage at this site, which agrees with analysis of the protein by SDS-PAGE (**Figure 1d**). The lack of density in the cryoEM maps is likely a result of this region’s weak attachment to the GP core domains, and as previously hypothesized by other groups^51,52^ likely explains the lack of glycan cap density in their antibody bound cryoEM structures. Additionally, as expected^52^, the GP2 HR2-stalk is resolved but is much more flexible in the cryoEM map and was therefore not built into the model. Only a single glycan is resolved in our cryoEM map at N563; other glycan sites are located on flexible peptides or domains that did not resolve (**Figure 1i**). Notably, N563 and N618 are two conserved N-linked glycan sites found in the GPs of all filoviruses^53^. Finally, our apo-state cryoEM structure of EBOV GP—determined at pH 7.5—has subtle deviations compared to apo state structures solved by X-ray crystallography at pH range spanning 5.0-5.2^14,32,47^, including within and adjacent to a hydrophobic pocket between GP1 and GP2 where several antiviral small molecule lead candidates bind^47,49,54–56^ (**Supplemental Figure 6**).

### GP-I53-50A Design, Production, and Characterization

We next generated constructs encoding EBOV-GP-Foldon as an N-terminal fusion to the trimeric nanoparticle component I53-50A via a short flexible Gly-Ser linker. The descriptive name for this construct is “EBOV-GP-I53-50A” (MDT-000868, **Figure 2a**). We initially produced and characterized this construct using small-scale lentiviral transduction in HEK293 cells^50^ and observed secretion of a uniform trimer that we purified by IMAC and SEC (**Supplemental Figure 7a,b**). Using the same construct design strategy EBOV, we next generated an I53-50A SUDV construct called SUDV-GP-I53-50A (MDT-000869, **Figure 2e**). SUDV-GP-I53-50A was also produced and characterized using small-scale lentiviral transduction in HEK293 cells^50^. The secreted SUDV-GP-I53-50A was found to be a reasonably uniform trimer after purification by IMAC and SEC (**Supplemental Figure 7c,d**). Negative stain EM characterization of both EBOV-GP-I53-50A and SUDV-GP-I53-50A trimers revealed the expected molecular envelope for the two-domain homotrimeric proteins (**Figures 2b,c,d for EBOV and 2f,g,h for SUDV, Supplemental Figure 13a**).

**Fig. 2:**
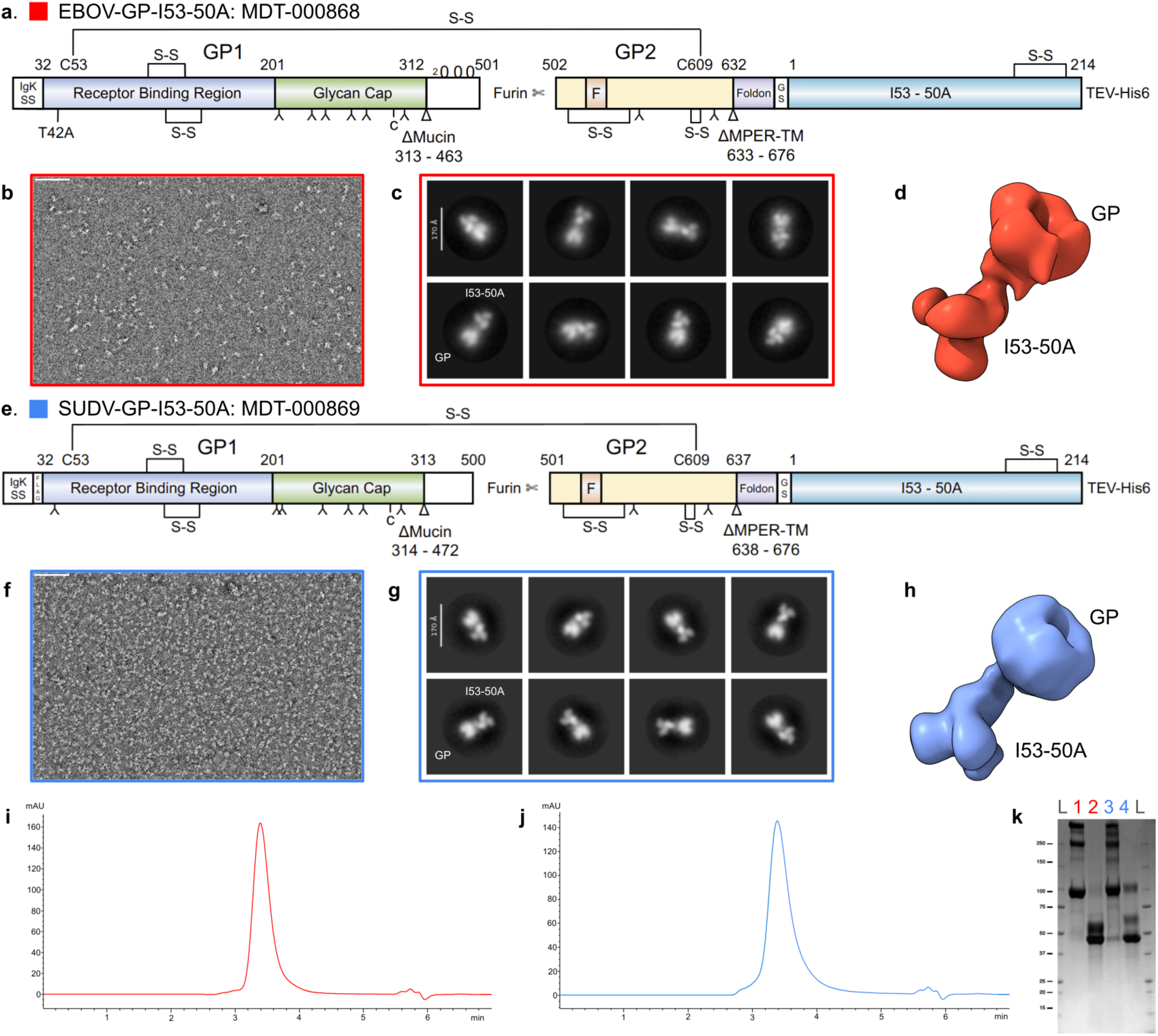
Production and Characterization of EBOV-GP-I53-50A and SUDV-GP-I53-50A Constructs. The domain architectures of EBOV-GP-I53-50A MDT-000868 (red) and SUDV-GP-I53-50A MDT-000869 (blue) using Figure 1a schema are illustrated in **a.** and **e.** Secreted proteins were expressed in HEK293 cells by lentiviral transduction and purified from serum free medium by IMAC and SEC and then characterized by negative stain electron microscopy (see Methods, and **Supplemental Figure 13a**) in **b.** and **f.** 2D-classification and 3D reconstructions of GPs were generated from negative stain EM micrographs for EBOV in **c.** and **d.** and SUDV in **g.** and **h.** EBOV-GP-I53-50A (MDT-000868) and SUDV-GP-I53-50A (MDT-000869) constructs were produced by transient transfection using pCMV/R vectors and purified from serum free medium by IMAC and SEC shown in **i.** and **j.** Final SEC pure proteins were characterized by analytical SEC HPLC in **i.** (EBOV, red) and **j.** (SUDV, blue) and **k.** SDS-PAGE under non-reducing (NR, lanes 1 and 3) and reducing (R, lanes 2 and 4) conditions prior to being used for nanoparticle assemblies and animal immunization studies. Molecular weight markers in ladder (L) are shown. See **Supplemental Figure 13a** for additional details on nsTEM of these NP immunogens.

### I53-50 Nanoparticle Immunogen Production and Structural Characterization

We then scaled up expression and purification of EBOV-GP-I53-50A and SUDV-GP-I53-50A by transient transfection of ExpiHEK293F cells, generating >10 milligrams of each after purification by IMAC and SEC (**Supplemental Figure 8**). The purified proteins had uniform analytical SEC-HPLC profiles and the expected SDS-PAGE band patterns (**Figure 2 i,j,k**). Following previously reported protocols^23^, we then mixed the GP-I53-50A trimers with pentameric I53-50B pentamer purified from *E. coli* to produce icosahedral NP immunogens displaying 20 GP trimers (illustrated in **Figure 3a**). We produced and characterized four types of GP-I53-50 NP immunogens in this way: EBOV-GP-I53-50 NP (**Supplemental Figure 9**), SUDV-GP-I53-50 NP (**Supplemental Figure 10**), a mosaic I53-50 NP co-displaying EBOV- and SUDV-GP in a 1:1 ratio (**Supplemental Figure 11**), and a 1:1 “cocktail” mixture of EBOV-GP-I53-50 + SUDV-GP-I53-50 (**Supplemental Figure 12**). Each of these was further structurally validated by nsEM, and the observed 2D class averages revealed the expected underlying I53-50 NP chassis and GP extensions (**Figure 3b,c,d**). Ab-initio and 3D refinement was performed for each sample, and the resulting maps suggested that each particle was well-formed (**Supplemental Figure 13b**).

**Fig. 3:**
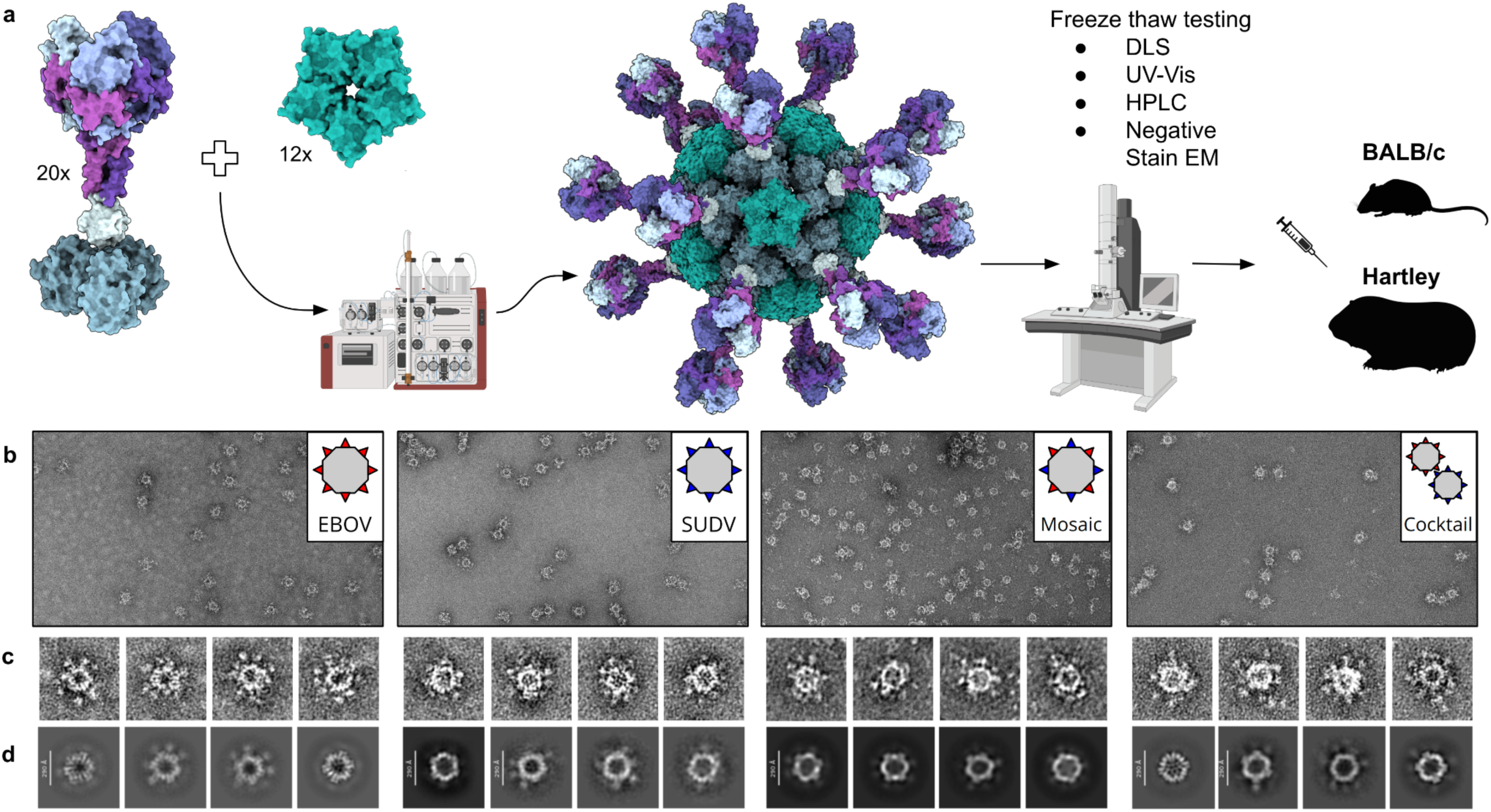
Production and Negative Stain EM Characterization of EBOV- and SUDV-GP-I53-50A Antigens as Assembled I53-50 Nanoparticles. **a.** The 3D molecular representations of the structure of EBOV- and SUDV-GP-I53-50A trimeric antigens along with the pentameric I53-50B NP component. When mixed in solution at approximately 1:1 stoichiometric ratios the proteins self-assemble into icosahedral two-component nanoparticles with molecular structure illustrated with the copy numbers noted. The assembled NP immunogens are purified by SEC. Peak fractions are aliquoted, flash frozen, and tested for particle uniformity pre and post freeze thaw by dynamic light scattering DLS, UV-Vis, analytical SEC HPLC, and negative stain TEM (see **Supplemental Figures 9-23, and 21-23**) prior to rodent immunization studies. Created with biorender.com and ChimeraX. **b. c. d.** are arranged in columns and rows with EBOV (red) and SUDV (blue) NP immunogens illustrated. **Row b.** The negative stain TEM micrographs. **Row c.** The 5Å Low-pass filtered single particle images extracted from micrograph. **Row d.** The 2D class averages generated from all particles in each dataset. The I53-50 icosahedral chassis are well resolved with limited visibility of flexible GP components. **Column 1**, EBOV-GP-I53-50 NP. **Column 2**, SUDV-GP-I53-50 NP. **Column 3**, Mosaic NP where EBOV-GP-I53-50A and SUDV-GP-I53-50A are co-assembled with 153-50B. **Column 4**, Cocktail of EBOV-GP-I53-50 NP and SUDV-GP-I53-50 NP mixed post purification of individual nanoparticles. See **Supplemental Figure 13b** for additional details on nsTEM of these NP immunogens.

### Prophylactic Immunization and Challenge with Mouse-adapted EBOV

Having confirmed the structural integrity of our EBOV- and SUDV-GP-I53-50 NP immunogens, we next conducted a prime-boost mouse immunization study including a challenge with maEBOV^57^ in the USAMRIID BSL-4 facilities (**Figure 4a**). The study included 8 groups (10 mice per group), and all test articles were formulated with an equal volume of AddaVax™, a squalene-based oil-in-water adjuvant, prior to subcutaneous injection. Four main test Groups 1-4 included EBOV-GP-I53-50 (Group 1), SUDV-GP-I53-50 (Group 2), the mosaic EBOV / SUDV-GP-I53-50 NP (Group 3), and the 1:1 cocktail of EBOV-GP-I53-50 and SUDV-GP-I53-50 NPs (Group 4). To investigate the importance of nanoparticle assembly in eliciting anti-GP antibody responses, we also included matched “non-assembling” control Groups 5-7 using the same doses of the GP-I53-50A trimers (2.5 µg) mixed with a pentameric protein called “2OBX” (0.8 µg)^58,59^ that is nearly identical to the I53-50B pentamer, but is not capable of driving assembly of the icosahedral I53-50 nanoparticle^22^, as it lacks the required computationally designed protein-protein interface with I53-50A (**Supplemental Figures 14-16**). As a negative control immunogen (Group 8), we also injected animals with bare I53-50 NP (2.5 µg) lacking any GP antigen (**Supplemental Figure 17**).

**Fig. 4:**
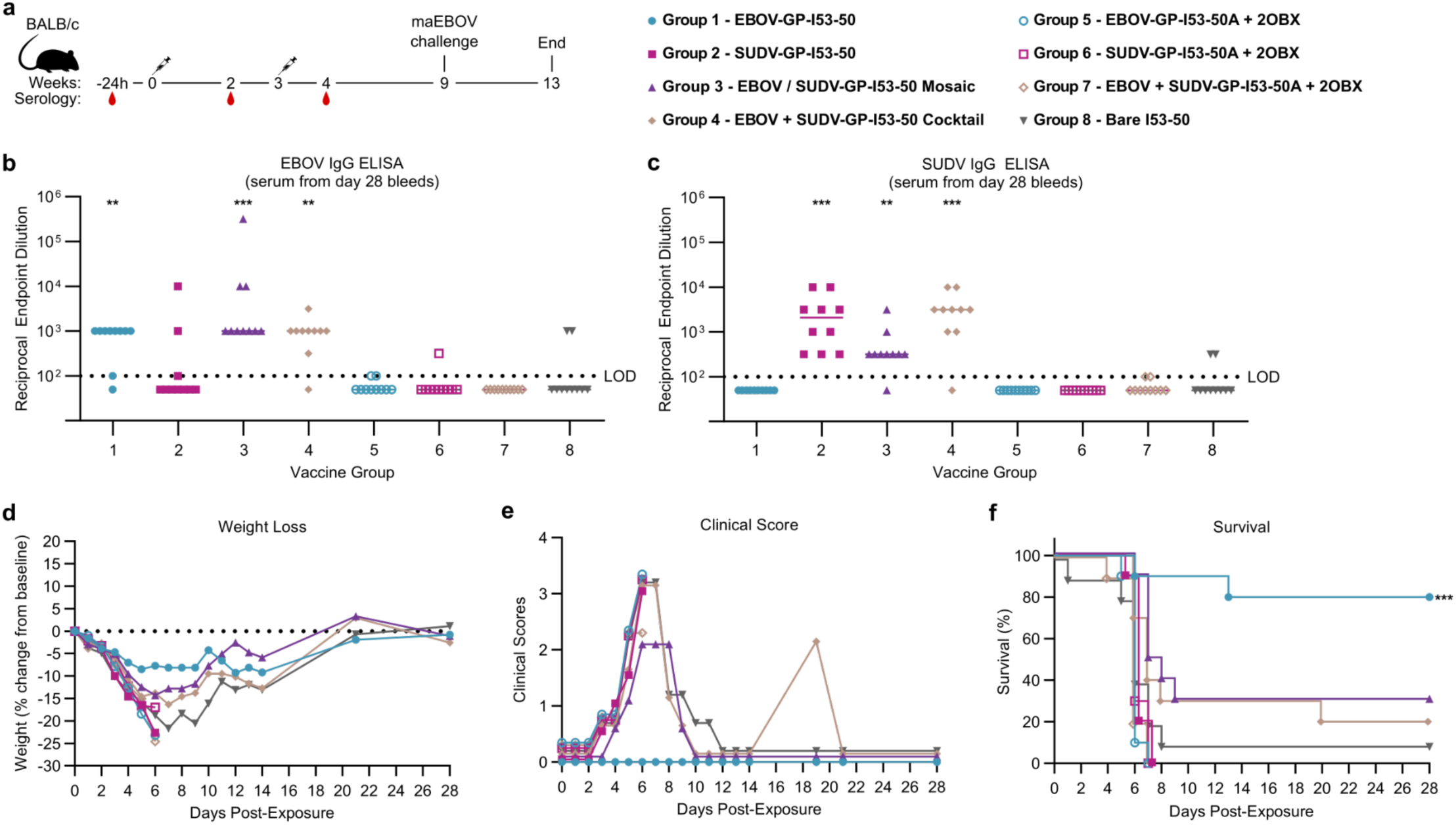
Prophylactic Immunization and Challenge with the Mouse Adapted EBOV. **a.** The study protocol is illustrated. Serum was collected from mice the day prior to a prime dose, two weeks following the prime dose, and one week following the boost dose at week 3. At week 9, animals were transferred to BSL-4 facilities and challenged with 100 pfu of maEBOV by intraperitoneal (IP) route after which animals were monitored for moribund condition and weight loss as a sign of EVD for 28 days following viral challenge. Test article descriptions are noted in the legend with colorized dots for each group (10 animals per group) **b.** Anti-EBOV GP or **c.** anti-SUDV GP IgG antibody titers from day 28 post-boost blood collections were measured by ELISA following serial dilution to establish the Y-Axis, Log10 plot of the reciprocal endpoint of dilution needed to observe signal in the assay relative to baseline pre-immunization serum signals. **d.** Average daily percentage weight loss or gain over time was measured for each group of animals following viral challenge. **e.** Average daily clinical disease scores were monitored for each group of animals following viral challenge. **f.** Kaplan–Meier survival curves, where moribund mice were promptly euthanized when they met predefined euthanasia criteria, a score of 4; defined as bleeding, unresponsive, weak, or unable to walk. Legend for Groups is indicated in the upper right. For survival, P values are indicated for comparisons against the respective bare I53-50 control Group 8, based on log-rank Mantel-Cox statistical analyses, *P < 0.05; **P < 0.01; ***P < 0.001.

Serological responses at day 28 (7 days after boost vaccination) demonstrated strong antibody titers to EBOV-GP in Group 1 or SUDV-GP in Group 2 (**Figure 4b,c**). Little or no cross-reactive antibody responses were observed: only 3 out of 10 animals that received EBOV-GP-I53-50 had any observable anti-SUDV-GP antibodies, while none of the ten animals that received SUDV-GP-I53-50 had any appreciable anti-EBOV-GP antibodies. This was expected as the EBOV and SUDV GP components of our NP immunogens have only 68% sequence identity (**Supplemental Figures 18 and 19**), and previous reports on filovirus vaccine candidates have suggested that little heterologous cross-protection can be obtained with single filoviral GP immunogens. Notably, animals immunized with either the mosaic (Group 3) or cocktail (Group 4) immunogens elicited strong antibody responses against both EBOV-GP and SUDV-GP, with only one or zero of the animals in Groups 3 or 4 lacking an immune response to either EBOV-GP or SUDV-GP. As evidence that antigen display in the form of a nanoparticle is important for eliciting potent immune responses, the non-assembling controls of EBOV-GP-I53-50A with 2OBX (Group 5), SUDV-GP-I53-50A with 2OBX (Group 6), or an equal mixture of EBOV- and SUDV-GP-I53-50A with 2BOX (Group 7) did not elicit strong antibody responses (e.g., compare Group 1 to 5 or Group 2 to 6). Sera from two of the Group 8 mice receiving the bare I53-50 nanoparticle test article unexpectedly appeared to have low but detectable anti-EBOV-GP and anti-SUDV-GP IgG titers.

To determine if the immunogens were capable of protecting animals from maEBOV challenge^60^, five weeks after the booster vaccination all animals were exposed to 100 plaque forming units (pfu) of maEBOV by intraperitoneal (IP) injection, and then monitored daily for signs of disease (morbidity is defined as having lost >15% of body weight, bleeding, unresponsive, weak, or unable to walk). Comparing the Kaplan–Meier survival curves in this challenge model indicated that a statistically significant (p < 0.001) protective advantage was afforded to the groups of animals receiving EBOV-GP-I53-50 (Group 1); these animals lost less weight and had better clinical scores and survival than all other groups (**Figures 4d-f**). Some of the mice receiving the mosaic EBOV/SUDV-GP-I53-50 NP (Group 3) or cocktail EBOV-GP-I53-50 + SUDV-GP-I53-50 (Group 4) were also protected from challenge, although the differences were not statistically significant compared to the negative control immunogen (bare I53-50 NP; Group 8). Poor survival was observed in mice receiving only SUDV-GP-I53-50 (Group 2) and the non-assembling control immunogens (Groups 5-7) where all animals succumbed by day 8 post-challenge. One Group 8 animal survived maEBOV challenge, but it was not one of the two that had detectable anti-EBOV antibodies (**Figure 4b**). Even though the mosaic (Group 3) and cocktail (Group 4) mice displayed statistically significant anti-EBOV GP or anti-SUDV GP IgG responses relative to bare I53-50 controls (**Figures 4b,c**), respectively only three or two of the mice in each of these groups survived maEBOV challenge. While this survival result is not statistically significantly better than the observed survival in control bare I53-50 Group 8 (which had one surviving animal), it is statistically significant when compared to the SUDV-GP-I53-50 vaccinated animals (Group 2) for mosaic (Group 3, p < 0.001) or cocktail (Group 4, p < 0.01) (**Supplemental Figure 20a**). Collectively, these results indicate that EBOV-GP displayed on the I53-50 NP elicited protective anti-EBOV-GP responses, whereas non-assembling GP-I53-50A trimers did not. By contrast, while the SUDV-GP-I53-50 nanoparticle immunogen was capable of eliciting anti-SUDV-GP IgG responses, this immune response was not cross-protective in mice subsequently challenged with maEBOV.

### Prophylactic Immunization and Challenge with the Guinea Pig-adapted SUDV

Having established that the EBOV-GP-I53-50 nanoparticle immunogen can elicit protective immune responses, we conducted a follow-up prophylactic prime-boost immunization study with Hartley guinea pigs, including challenge with guinea pig-adapted SUDV (gpaSUDV)^61^ in the USAMRIID BSL-4 facilities (**Figure 5**). Groups of 6 animals (except for Group 1 which had 5 animals) were immunized with newly prepared nanoparticle immunogens similar to those we used in the mouse study, receiving 2.1 µg of GP antigen in each injection (**Supplemental Figures 21-23**). We did not include the non-assembling 2OBX groups. As a negative control immunogen, we injected animals with 2.9 µg of bare I53-50 NP lacking any GP antigen (Group 5, see **Supplemental Figure 17** for an example of this nanomaterial). All test articles were formulated with an equal volume of AddaVax™, prior to subcutaneous injection.

**Fig. 5:**
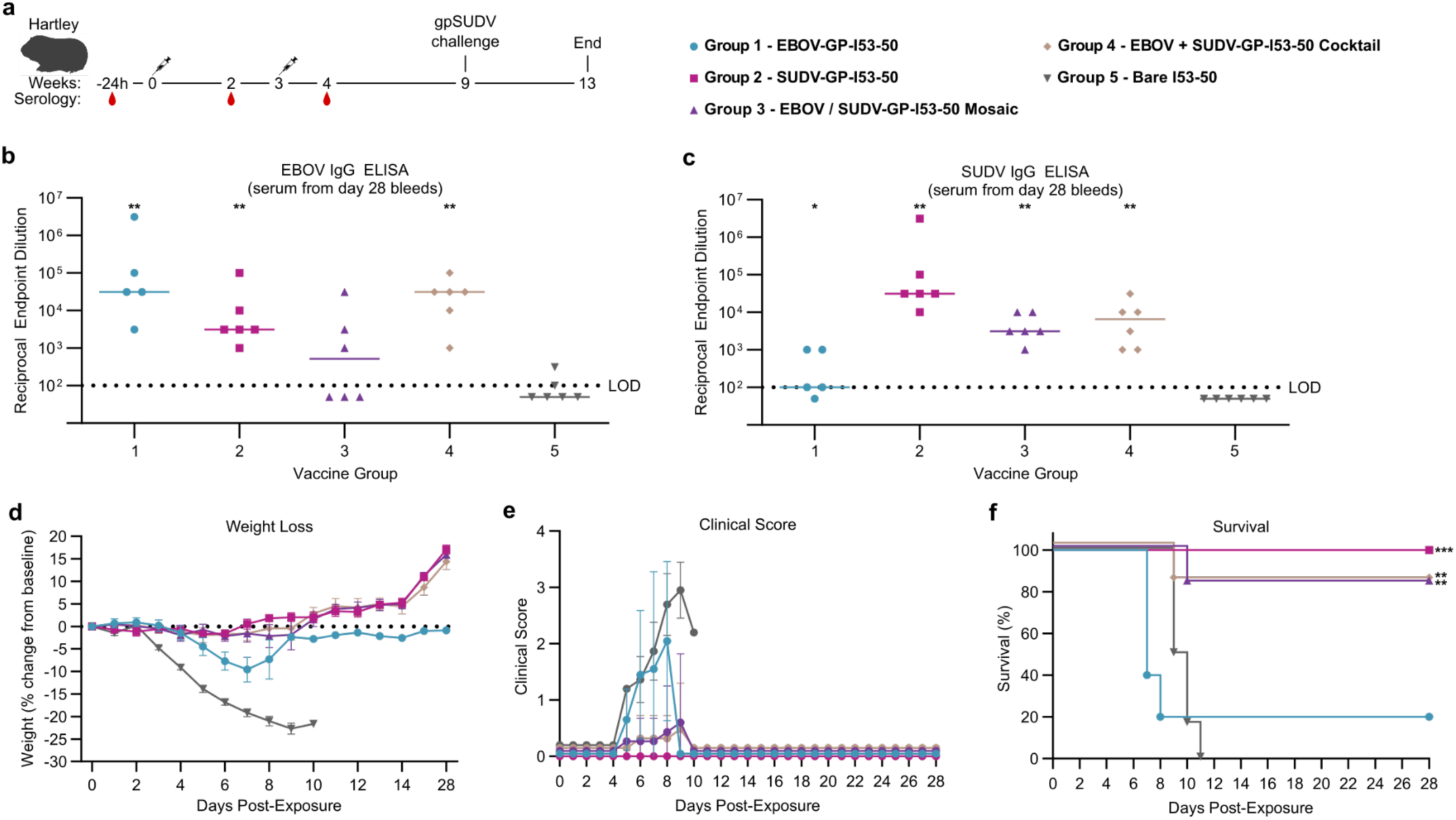
Prophylactic Immunization and Challenge with the Guinea Pig Adapted SUDV. **a.** The study protocol is illustrated. Serum was collected from mice the day prior to a prime dose, two weeks following the prime dose, and one week following the boost dose at week 3. At week 9, animals were transferred to BSL-4 facilities and challenged with 100 pfu of gpaSUDV by IP route after which animals were monitored for weight loss as a sign of EVD for 28 days following viral challenge. Test article descriptions are noted in the legend with colorized dots for each group (6 animals per group, except for Group 1 which had 5 animals) **b.** Anti-EBOV GP or **c.** anti-SUDV GP IgG antibody titers from day 28 post-boost blood collections were measured by ELISA following serial dilution to establish the Y-Axis, Log10 plot of the reciprocal endpoint of dilution needed to observe signal in the assay relative to baseline pre-immunization serum signals. **d.** Average daily percentage weight loss or gain over time following challenge (weights for each animal were measured daily). **e.** Average daily clinical disease scores over time following viral challenge (clinical scores for each animal were measured daily). **f.** Kaplan– Meier survival curves, where moribund guinea pigs were promptly euthanized when they met predefined euthanasia criteria, a score of 4; defined as bleeding, unresponsive, weak, or unable to walk. Legend for Groups is indicated in the upper right. For survival, P values are indicated for comparisons against the respective bare I53-50 control Group 5, based on log-rank Mantel-Cox statistical analyses, *P < 0.05; **P < 0.01; ***P < 0.001.

As observed in the previous mouse study, serological responses in guinea pigs at 7 days post boost (day 28) demonstrated strong vaccine-matched IgG antibody titers to EBOV-GP in Group 1 or SUDV-GP in Group 2 (**Figure 5a-c**). EBOV-GP-I53-50 did not generate cross-reactive IgG antibody responses to SUDV-GP, just as we observed in mice (**Figure 4c**, **Figure 5c**). However, the SUDV-GP-I53-50 NP did produce detectable cross-reactive antibody responses to EBOV-GP in guinea pigs, a result that we did not observe in mice (**Figure 4b**, **Figure 5b**) but has been observed in previous vaccine studies using mixtures of human parainfluenza virus type 3 (HPIV3) vectors expressing EBOV, SUDV, and BDBV GPs (see Figure 2 in ^62^). The mosaic Group 3 and cocktail Group 4 immunogens both elicited anti-GP IgG responses to EBOV-GP and SUDV-GP, although anti-EBOV-GP IgG titers were only detectable in three of the Group 3 animals. None of the Group 5 animals receiving the bare I53-50 negative control had any appreciable anti-GP IgG.

At five weeks following the booster immunization, all animals were transferred to BSL-4 and dosed with IP injections of 1,000 pfu of gpaSUDV. Animals immunized with any I53-50 NP bearing the SUDV-GP antigen (Groups 2-4) had an 80-100% survival rate at 28 days post-infection (**Figures 5d-f**). Furthermore, these animals remained asymptomatic or demonstrated mild symptoms as measured by temperature (**Supplemental Figure 24**) and overall clinical disease score (**Figures 5e**). By contrast, animals immunized with EBOV-GP-I53-50 (Group 1) or the bare I53-50 NP (Group 5) had limited survival rates and showed high levels of clinical disease. All of the Group 5 control animals succumbed by day 11, and only one of the guinea pigs in Group 1 recovered 9 days after infection. Comparing Kaplan–Meier survival curves indicated that a statistically significant protective advantage was afforded to the animals receiving SUDV-GP-I53-50 (Group 2; p < 0.001), the mosaic EBOV/SUDV-GP-I53-50 NP (Group 3; p < 0.01), or the EBOV-GP-I53-50 + SUDV-GP-I53-50 NP cocktail (Group 4; p < 0.01) compared to the control group treated with bare I53-50 (Group 5). Even though the mosaic (Group 3) and cocktail (Group 4) guinea pigs displayed either an average or statistically significantly higher anti-EBOV GP or anti-SUDV GP IgG response relative to bare I53-50 controls (**Figures 5b,c**), five of the six guinea pigs in each of these groups respectively survived gpaSUDV challenge, a significantly better survival for mosaic (Group 3, p < 0.05) or cocktail (Group 4, p < 0.01) when compared to the EBOV-GP-I53-50 vaccinated animals (Group 1) (**Supplemental Figure 20b**). Overall, these survival results were comparable to those of the maEBOV challenge, wherein statistically significant animal adapted virus protection can be achieved by either a mosaic or cocktail mix of EBOV- and SUDV-GP-I53-50 NP immunogens when compared to the heterologous GP-I53-50 groups; Group 2 SUDV-GP-I53-50 in the maEBOV study, or Group 1 EBOV-GP-I53-50 in the gpaSUDV study (**Supplemental Figure 20**).

## Discussion

A growing body of prophylactic vaccination data from both animal models and in people have shown that vaccines with only EBOV-GP or SUDV-GP immunogens alone are not capable of eliciting cross-Ebolavirus anti-GP immune responses and therefore do not offer broad-spectrum EVD protection^14,15^. Nevertheless, GP has been a productive target of monoclonal monoclonal antibody (mAb) cocktails to achieve pan-acting protection against EBOV and SUDV. These include MBP134^AF^ (ADI-15878^AF^ + ADI-23774^AF^)^63^, CA45 and FVM04^64^, as well as broadly neutralizing mAbs 4F9 and 6H8, each of which on its own provided protection against at least one Ebolavirus in relevant animal models, while the cocktail of both mAbs conferred superior protection against SUDV in guinea pigs^65^. Studies on structural determinants of the mAb-GP interactions for FDA-approved EVD mAb therapies including Ebanga™ and Inmazeb™, and ZMapp™(ref. ^66^) mAb cocktails for treatment of SUDV infection^67^ as well as the identification of single pan-filovirus mAbs^68^ have all pointed to the need to target multiple distinct epitopes on Ebolavirus GPs^52^. As such, in this study we sought to design and test the efficacy of I53-50 NP vaccines displaying either EBOV-GP or SUDV-GP, as well as mosaic and cocktail versions of the NP immunogens.

We have demonstrated that I53-50 protein nanoparticles co-displaying prefusion GP (Δmuc, ΔMPER-TM) antigens from Zaire Ebolavirus and Sudan Ebolavirus can provide prophylactic protection against death and disease symptoms (e.g., clinical disease score and weight loss) in two separate BSL-4 animal models (maEBOV mouse and gpaSUDV guinea pig models). Furthermore, I53-50 nanoparticles with individual EBOV-GP or SUDV-GP antigens demonstrated increased anti-GP IgG serum responses and protection from death and weight loss in the maEBOV and gpaSUDV rodent models. In our maEBOV study, we compared the antibody responses and protection provided by the GP-I53-50 NP immunogens to identical quantities of soluble trimeric GP-I53-50A mixed with a non-assembling 2OBX pentamer. The weak antibody responses in the groups of mice receiving the non-assembling 2OBX immunogens demonstrate that NP display of the GP antigen is key to eliciting a strong and protective antiviral immune response.

As part of our immunogen design work, we also provide the first high-resolution cryo-EM structure (3.05 Å) of prefusion trimeric EBOV-GP without any other binding partners, such as antibody fragments, which provides new insights to help support antiviral protein minibinder design or small molecule drug discovery. While previous EBOV GP structures have been solved in the apo state by X-ray crystallography^14,32,47^, all reported structures were generated under similar crystallization conditions. These include the use of similar buffers (PEG, sodium citrate), crystal packing in space groups H32 and P321 (with a shared crystal packing interface near inhibitor binding site), and under a pH range spanning 5.0-5.2 (**Supplemental Figure 6a**)^14,32,47^. Importantly, crystal packing in these structures consistently forms in close proximity to a small-molecule binding site, which consists of a hydrophobic pocket between the GP1 and GP2 subunits. In apo structures, this pocket is occupied by F193 and F194 which are displaced upon binding of several small molecule antiviral lead candidates that have been shown to inhibit viral entry and make GP susceptible to thermal destabilization^47,49,54–56^. In the analysis of our apo-state cryoEM structure of EBOV GP—determined at pH 7.5—we highlight notable deviations from prior crystal structures in this pocket and nearby segments that are involved in crystal contacts of reported apo-forms of GP (**Supplemental Figure 6**). These differences reflect the solution-state conformation of the EBOV GP small-molecule binding pocket under physiological conditions, which could have important implications for future rational development of additional small-molecule antivirals targeting EBOV. Moreover, this structural information could also guide the development of stabilized Ebolavirus GP antigens for next-generation vaccines or antibody design.

There are several limitations to the studies reported here. Our serological analyses of anti-GP IgG responses were based purely on ELISA assays, as the blood volumes and BSL-4 facility expenses required to assess anti-viral neutralization prevented such studies. Furthermore, we have not examined the epitope specificities or structures of monoclonal or polyclonal antibodies elicited by our NP immunogens, and therefore do not know if similar antibodies are induced by the mosaic and cocktail versions of our GP-I53-50 NP immunogens. Notably, co-display of multiple SARS-CoV-2 spike RBDs viral antigens in the I53-50 chassis can elicit neutralizing antibodies against distinct variants of concern (VOCs)^35,69^. Future studies could potentially explore whether the inclusion of additional Ebolavirus GPs, such as those from Marburg virus (MARV), Bundibugyo virus (BDBV), or Taï Forest virus (TAFV) could provide protective immunity against an even broader spectrum of EVD-causing filoviruses. In addition, our NP immunogens lacked the MPER region of GP, which is known to be targeted by broadly neutralizing antibodies^70–72^. Although display of an MPER-based epitope scaffold on the computationally designed two-component I53-40 nanoparticle failed to elicit neutralizing antibodies after rabbit immunization^73^, inclusion of the MPER domains in prefusion-stabilized trimeric GPs displayed on nanoparticles may further improve vaccine-elicited antibody responses. Future research would also benefit from an exploration of the nature of T cell responses to our GP-I53-50 NP immunogens^19^.

The two-component I53-50 NP chassis is well validated, and has already been licensed in multiple countries as a safe and effective vaccine technology, displaying 60 copies of the SARS-CoV-2 spike RBD for COVID-19 ^20,21,74^ - SKYCovione™. The I53-50 NP chassis has a growing safety profile with multiple clinical trials for SKYCovione™ and RSV/HMPV^34^ demonstrating that scaled manufacturing of I53-50 enabled vaccines is not likely to present a roadblock to further clinical development of the EBOV- and SUDV-GP-I53-50 vaccines presented here. Notably, in our hands even at this early stage of development, the production EBOV-GP-I53-50A and SUDV-GP-I53-50A components was respectively only ∼four-fold lower (∼7.3 or 7.6 mg/L by transient transfection (**Supplemental Figure 8**) than that observed for SARS-CoV-2 RBD-I53-50A fusion proteins^19^. This suggests that the filoviral immunogens reported here have a strong probability of clinical developability. In comparison, single-component nanoparticle vaccine candidates based on I3-01^32,58^ or ferritin^30^ cannot be used to produce mosaic nanoparticles displaying oligomeric antigens, which our results suggest may be needed to elicit broad-spectrum immune responses. Mosaic nanoparticles can be produced using the SpyTag/SpyCatcher system^75,76^, but this approach has yet to be de-risked at scale.

By following a structured EM-based characterization pipeline, the EBOV-GP and SUDV-GP trimeric antigens used in this study were identified as properly folded and assembled as trimeric antigens, remained well-folded when fused to the I53-50A nanoparticle component, and retained their integrity upon final assembly into vaccine nanoparticles. This structural validation translated to protective efficacy in two independent rodent studies. Collectively, these results support further development of the EBOV/SUDV-GP-I53-50 immunogen as a pan-EBOV-SUDV prophylactic vaccine candidate, bringing us one step closer toward a universal filovirus vaccine that provides protection against multiple filovirus species. As has been suggested by other EBOV-GP nanoparticle vaccine studies^32^, there may be value in boosting people who have already received the FDA-approved Ervebo^®^ vaccine with an all-protein GP nanoparticle immunogen vaccine to provide extended protection from EVD. Thus, the I53-50 nanoparticle platform for EBOV and SUDV may be complementary in clinical use to the rVSV vaccines for these two EVD-causing filoviruses.

## Methods

### Production of EBOV and SUDV Glycoprotein Immunogens by Lentiviral Transduction

MDT-000253, MDT-000868, MDT-000869, and MDT-000759 constructs (see detailed sequences in **Table 1**) were expressed as a soluble secreted proteins by lentiviral transduction using the Daedalus expression method^50^ using HEK293F (Invitrogen) cells grown in the presence of kifunensine to limit N-glycan heterogeneity. The secreted proteins were captured from conditioned media using a HisTrap FF crude column and eluted in PBS containing 250 mM imidazole. Each protein was then polished using a Superdex 200 Increase 10/300 GL column on an AKTA Pure M (GE) using a phosphate buffered saline (PBS) mobile phase. SDS-PAGE analysis was performed using NuPAGE 4-12% Bis-Tris gels run under non-reducing or reducing conditions where samples are treated with 1 mM dithiothreitol (DTT) prior to electrophoresis using manufacturer’s protocols. Final production yields were calculated based on absorbance determined by measuring absorbance at 280 nm using a UV-Vis spectrophotometer (Agilent Cary 8454) and calculated extinction coefficients^77^.

**Table 1.**
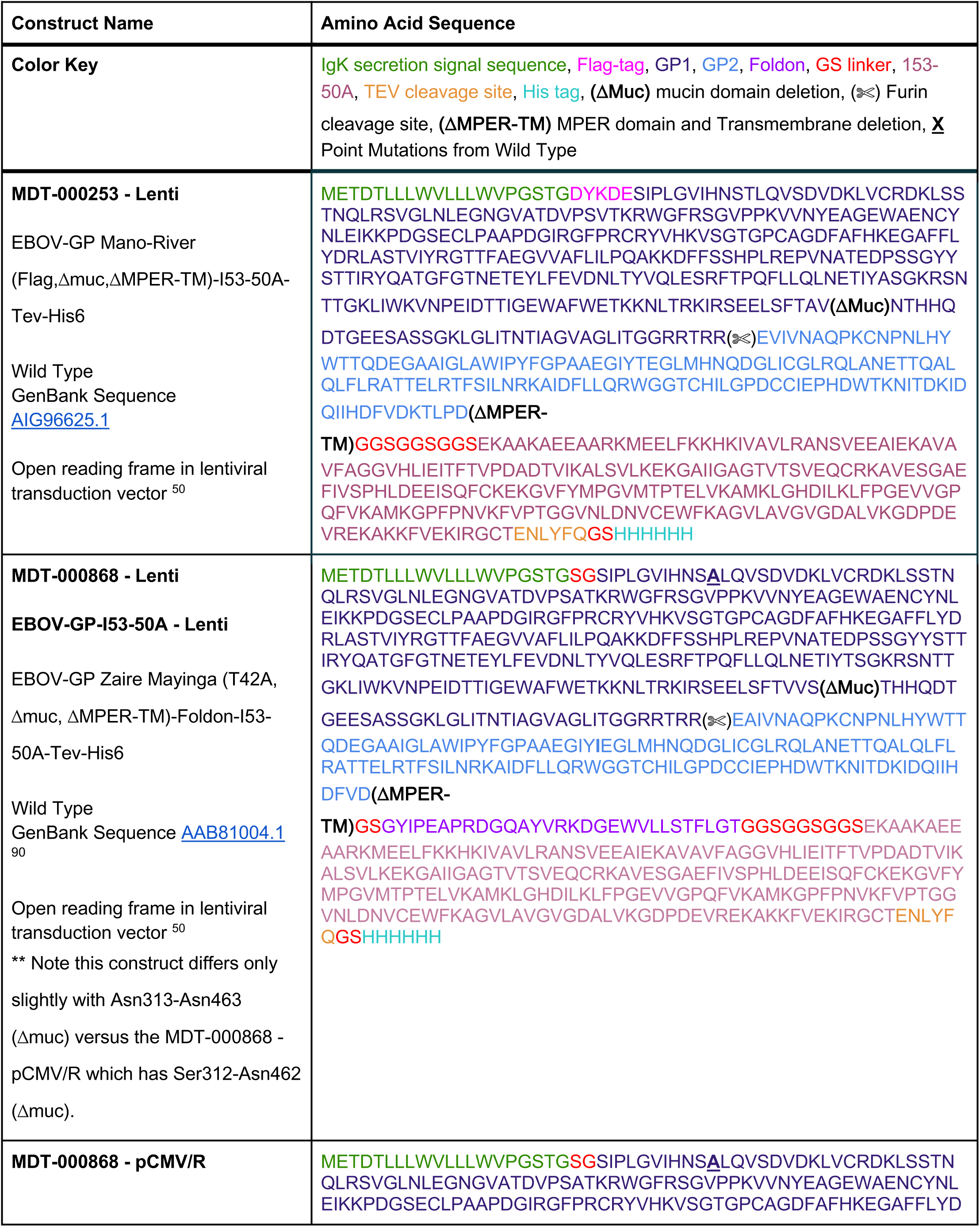
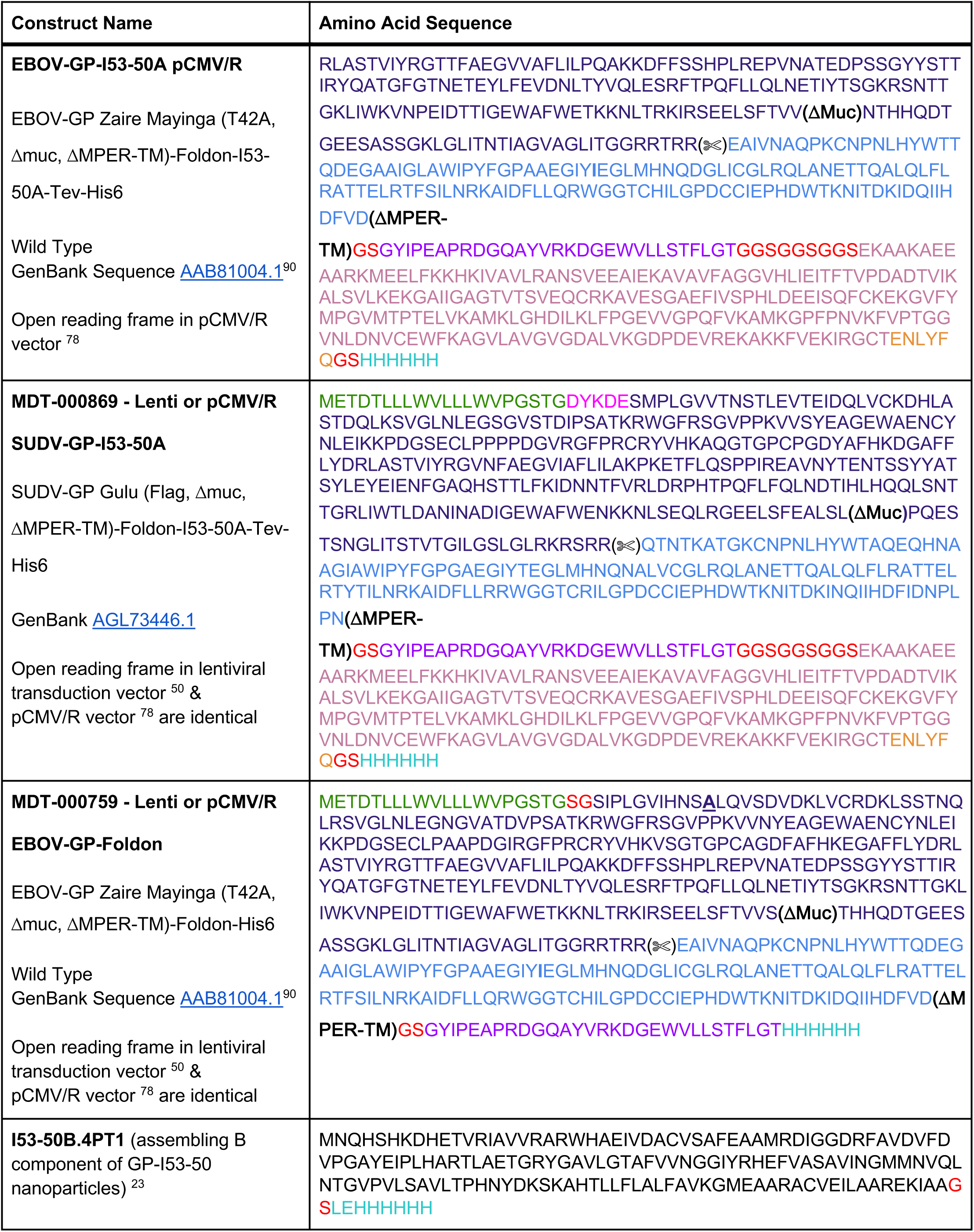
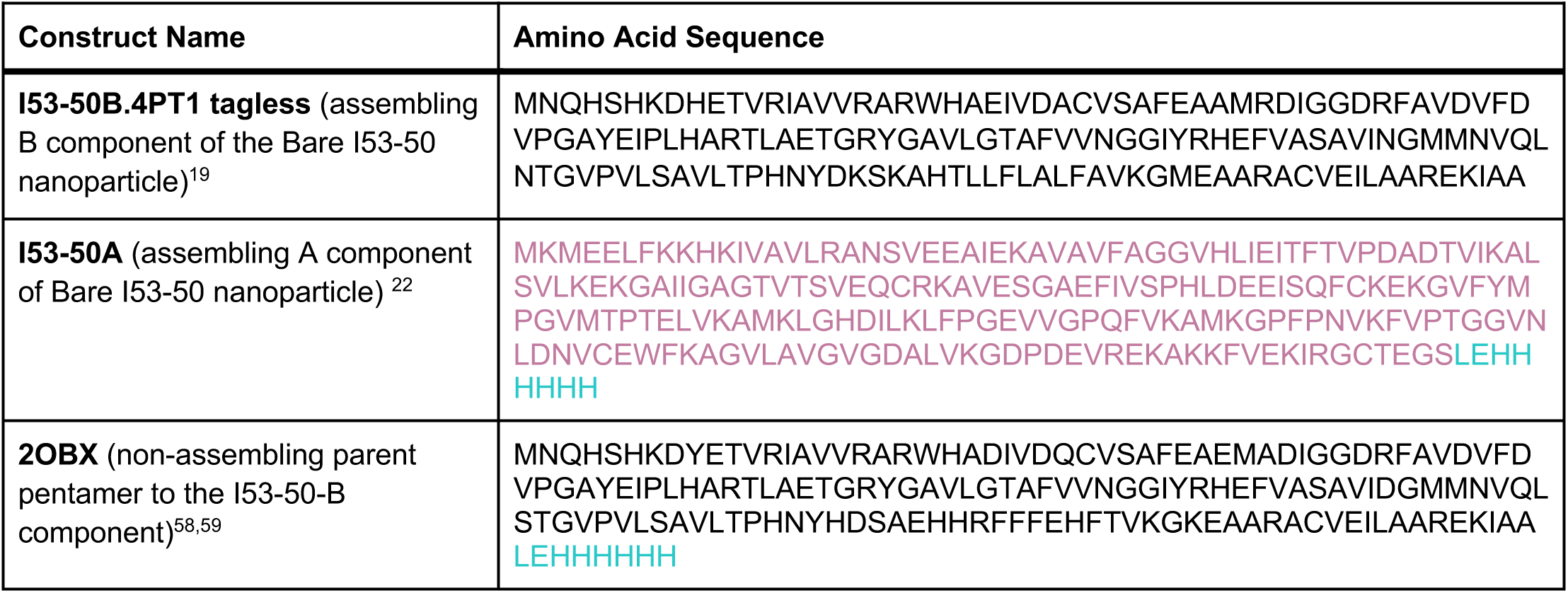
Sequences of EBOV and SUDV Glycoprotein Expression Constructs. The table below describes the exact open reading frames of the constructs used in the presented research (See **Supplemental Figure 19** a sequence comparison of the GP open reading frames of these constructs).

### Production of EBOV and SUDV Glycoprotein Immunogens by Transient Transfection

Proteins were produced with with pCMV/R vectors^78^ with previously described methods^69^ using endotoxin-free DNA in ExpiHEK293F cells grown in suspension using Expi293F expression medium (Life Technologies) at 33°C, 70% humidity, 8% CO2 rotating at 150 rpm. The cultures were transfected using PEI-MAX (Polyscience) with cells grown to a density of 3.0 million cells per mL and cultivated for 3 days. Supernatants were clarified by centrifugation (5 minutes at 4000 rcf), addition of PDADMAC solution to a final concentration of 0.0375% (Sigma Aldrich, #409014), and a second centrifugation (5 minutes at 4000 rcf). Similar methods as noted above for purification of constructs by IMAC and SEC were applied to the purification of GP immunogens from culture medium following the transient transfections. Final production yields were calculated based on absorbance determined by measuring absorbance at 280 nm using a UV-Vis spectrophotometer (Agilent Cary 8454) and calculated extinction coefficients^77^.

### Microbial plasmid construction, protein expression and purification of Tagless I53-50B.4PT1 Tagless

I53-50B.4PT1 plasmid^23^ was synthesized by GenScript in pET29b between the NdeI and XhoI restriction sites with a double-stop codon. Protein was expressed in Lemo21(DE3) cells (NEB) in LB (10 g Tryptone, 5 g Yeast Extract, 10 g NaCl) grown in a 10 L BioFlo 320 Fermenter (Eppendorf). At inoculation, impeller speed was set to 225 rpm, sparge per liter per minute (SPLM) set to 5 with O2 supplementation as part of the dissolved-oxygen aeration cascade, and the temperature set to 37°C. At the onset of a dissolved oxygen (DO) spike (OD600 ∼12), the culture was supplemented with a bolus addition of 100 mL of 100% glycerol and induced with 1 mM IPTG. During this time, the culture temperature was reduced to 18°C, and O2 supplementation was ceased, with expression continuing until an OD600 ∼20. The culture was harvested by centrifugation and the pellets were resuspended in PBS, homogenized, and then lysed by microfluidization using a Microfluidics M110P at 18,000 psi using 3 discrete passes. Following sample clarification by centrifugation (24,000 *g* for 30 minutes), the supernatant was discarded, and protein was extracted from the inclusion bodies. First, the pellet was washed with PBS supplemented with 0.1% Triton X-100, pH 8.0. After this initial wash and sample clarification by centrifugation, the pellet was washed with PBS supplemented with 1 M NaCl, pH 8.0. Following the second wash, the protein was extracted from the pellet into PBS supplemented with 2 M urea and 0.75% CHAPS (3-[(3-Cholamidopropyl)dimethylammonio]-1-propanesulfonate), pH 8.0. Following extraction, the sample was applied to a DEAE Sepharose FF column (Cytiva) on an ӒKTA Avant150 FPLC system (Cytiva). After sample binding, the column was washed with 5 column volumes (CV) of PBS supplemented with 0.1% Triton X-100, pH 8.0, followed by a 5 CV wash with PBS supplemented with 0.75% CHAPs, pH 8.0 in series. The protein was eluted with 3 CV of PBS supplemented with 500 mM NaCl, pH 8.0. After purification, fractions were pooled and concentrated in 10K molecular weight cutoff (MWCO) centrifugal filters (Millipore), sterile-filtered (0.22 μm), and tested to confirm low endotoxin levels before use in nanoparticle assembly.

### Microbial plasmid construction, protein expression and purification His6 Tagged I53-50B.4PT1

The gene for I53-50B.4PT1 with a C-terminal hexa-histidine affinity tag was synthesized by GenScript in pET29b between the NdeI and XhoI restriction sites with a double-stop codon, and its microbial production and purification was identical to that of the 2OBX non-assembling pentamer.

### Microbial plasmid construction, protein expression and purification 2OBX Non-Assembling Pentamer

2obx plasmid was synthesized by GenScript in pET29b between the NdeI and XhoI restriction sites in frame of the C-terminal polyhistidine tag^22^. The 2OBX^59^ pentamer was expressed in Lemo21(DE3) (NEB) in LB (10 g Tryptone, 5 g Yeast Extract, 10 g NaCl) grown in 2 L baffled shake flasks. Cells were grown at 37°C to an OD600 ∼0.8, and then induced with 1 mM IPTG. Expression temperature was reduced to 16°C and the cells were shaken for ∼16 hours. The cells were harvested and lysed by microfluidization using a Microfluidics M110P at 18,000 psi in 25 mM Tris, 500 mM NaCl, 30 mM imidazole, 1 mM PMSF, 0.01 mg/mL DNAse, and 0.75% CHAPS. Lysates were clarified by centrifugation at 24,000 g for 30 minutes and applied to 5 mL of NiNTA resin (Qiagen) for purification by gravity IMAC. Protein of interest was eluted using 25 mM Tris pH 8, 500 mM NaCl, 0.75% CHAPS, 500 mM imidazole buffer. The elution was pooled, concentrated in 10K MWCO centrifugal filters (Millipore), sterile filtered (0.22 μm) and applied to a Superdex 200 Increase 10/300 using 25 mM Tris pH 8, 500 mM NaCl, 0.75% CHAPS buffer. After sizing, the component was tested to confirm low levels of endotoxin before using it for animal vaccination studies as part of test articles.

### Microbial plasmid construction, protein expression and purification of I53-50A

The I53-50A plasmid was synthesized by GenScript in pET29b between the NdeI and XhoI restriction sites in frame of the C-terminal polyhistidine tag^22^. The I53-50A trimer was expressed as described^22^. Cells were lysed by sonication (2.5 minutes total sonicating time in 2 second pulses) in 50 mM Tris pH 8, 500 mM NaCl, 20 mM Imidazole, 0.75% 3-[(3-Cholamidopropyl)dimethylammonio]-1-propanesulfonate (CHAPS), 0.1 mg/mL lysozyme, 0.05 mg/mL DNase, 0.05 mg/mL RNase, 5 mM MgCl2 and 1 mM phenylmethylsulfonyl fluoride (PMSF). Lysate was clarified by centrifugation at 33,000 × g for 20 minutes. Lysate supernatants were applied to HisTrap HP or FF columns (GE Healthcare) for purification by immobilized metal affinity chromatography (IMAC) on an AKTA Pure FPLC system (GE Healthcare). Protein of interest was eluted over a linear gradient of 20 mM to 500 mM imidazole in a background of 50 mM Tris pH 8, 500 mM NaCl, and 0.75% CHAPS buffer after washing with ∼10 column volumes wash buffer (elution buffer with 20 mM imidazole). Peak fractions were pooled, concentrated in 10K MWCO centrifugal filters, sterile filtered (0.22 μm) and applied to a Superdex 200 Increase 10/300 GL SEC column (GE Healthcare) using 25 mM Tris pH 8, 500 mM NaCl, 0.75% CHAPS buffer, 1 mM DTT. I53-50A elutes at ∼15-17 mL.

### Assembly Production and Purification of EBOV and SUDV Glycoprotein I53-50 Nanoparticles

Total protein concentration of purified individual nanoparticle components was determined by measuring absorbance at 280 nm using a UV/vis spectrophotometer (Agilent Cary 8454) and calculated extinction coefficients ^77^. The assembly steps were performed at room temperature with addition in the following order: EBOV-GP-I53-50A or SUDV-GP-I53-50A trimeric glycoprotein, followed by quality sufficient (q.s.) volume filling with buffer as needed to achieve desired final concentration, and finally either tagless or hexa-histidine tagged I53-50B.4PT1 pentameric component (in 50 mM Tris pH 8, 500 mM NaCl, 0.75% w/v CHAPS), with a molar ratio of GP-I53-50A: I53-50B.4PT1 of 1.1:1. q.s. buffer contained 50 mM Tris pH 8, 150 mM NaCl, 5% v/v glycerol. A Superose 6 Increase 10/300 GL column was used for nanoparticle purification with running buffer as 25 mM Tris pH 8, 150 mM NaCl, 5% v/v glycerol for NPs prepared for the maEBOV study (**Figure 4**) or 50 mM Tris pH 8, 150 mM NaCl, 5% glycerol, 100 mM L-arginine for NPs prepared for the gpaSUDV study (**Figure 5**). Assembled nanoparticles were sterile-filtered (0.22 μm) immediately prior to column application and following pooling of SEC fractions. The EBOV-GP-I53-50 and SUDV-GP-I53-50 were prepared as independent productions on at least three separate occasions. See **Supplemental Figures 9-17** for certificate of testing information on the GP nanoparticle immunogens prepared in November of 2019 with hexa-histidine tagged I53-50B.4PT1 for use in the maEBOV study (**Figure 4**). See **Supplementary Figures 21-23** for certificate of testing information on the GP nanoparticle immunogens prepared with tagless I53-50B.4PT1 in July of 2021 for use in the gpaSUDV study (**Figure 5**). The third preparations of these GP nanoparticle immunogens occurred in August of 2022 with tagless I53-50B.4PT1 (see **Supplemental Figure 13b and Figure 3** for nsTEM of the EBOV-GP-I53-50 prepared in August of 2022).

### Assembly Production and Purification of Bare I53-50 Nanoparticles

Bare I53-50 nanoparticles were produced as previously described^69^ using I53-50A and I53-50B.4PT1 with hexa-histidine tag for the maEBOV study (**Supplemental Figure 17**) or tagless I53-50B.4PT1 for the gpaSUDV study. Total protein concentration of purified individual nanoparticle components was determined by measuring absorbance at 280 nm using a UV/vis spectrophotometer (Agilent Cary 8454) and calculated extinction coefficients. The assembly steps were performed at room temperature with addition in the following order: I53-50A trimeric protein, followed by additional buffer (50 mM Tris pH 7.4, 185 mM NaCl, 100 mM L-arginine, 4.5% glycerol, and 0.75% w/v CHAPS) as needed to achieve desired final concentration, and finally I53-50B.4PT1 pentameric component (in 50 mM Tris pH 8, 500 mM NaCl, 0.75% w/v CHAPS), with a molar ratio of I53-50A:I53-50B.4PT1 of 1.1:1. All I53-50 in vitro assemblies were incubated briefly at room temperature before subsequent purification by SEC in order to remove residual unassembled I53-50A components. A Superose 6 Increase 10/300 GL column was used for nanoparticle purification. Assembled nanoparticles were purified in 50 mM Tris pH 7.4, 185 mM NaCl, 100 mM Arginine, 4.5% v/v glycerol, and 0.75% w/v CHAPS, and elute at ∼11 mL on the Superose 6 column. Assembled nanoparticles were sterile-filtered (0.22 μm) immediately prior to column application and following pooling of fractions. The naked I53-50 nanoparticle chassis samples have been produced and purified on numerous occasions^19,22,23,25,26,34^.

### Endotoxin Measurements

Endotoxin levels in protein samples were measured using the EndoSafe Nexgen-MCS System (Charles River). Samples were diluted 1:100 in Endotoxin-free LAL reagent water, and applied into wells of an EndoSafe LAL reagent cartridge. Charles River EndoScan-V software was used to analyze endotoxin content, automatically back-calculating for the dilution factor. Endotoxin values were reported as EU/mL which were then converted to EU/mg based on UV/vis measurements. Our threshold for samples suitable for immunization was <50 EU/mg.

### CryoEM Sample Preparation, Data Collection, and Data Processing for EBOV-GP-Foldon

To prepare the samples, 2 μL of EBOV-GP-Foldon produced from our pCMV/R vector by transient transfection and purified by IMAC and SEC at a concentration of 1 mg/mL in 300 mM NaCl, 20 mM Tris, pH 7.5, was applied to glow-discharged, glow discharged for 25 seconds at 15 mA on 2.0/2.0-T C-flat holey carbon grids. Vitrification was performed using a Mark IV Vitrobot at 22 °C with 100% humidity for all. Blotting was done using a 7.0 second blot time, a blot force of 0 and a 7.5 second wait time was used before being immediately plunge frozen into liquid ethane. The CryoEM dataset for EBOV-GP-Foldon was collected automatically using Leginon^79^ and used to control a ThermoFisher Titan Krios 300 kV, microscopes was equipped with a K3 Summit direct electron detector^80^ and BioQuantum Gif energy filter and operated in counting mode. Random defocus ranges spanned between −0.8 and −1.8 μm using image shift. 3,128 movies with a pixel size of 0.84, total dose of 56.7 e^-^/Å^2^ was recorded with a total exposure of 5 seconds over 100 frames.

All data processing was carried out in CryoSPARC^81^. Default parameters were used for refinement unless otherwise noted. The video frames were aligned using Patch Motion, and defocus and astigmatism values were estimated using the Patch Contrast Transfer Function. Curate Exposures was used to remove 1007 micrographs, leaving 2,121 good micrographs. In total, 1,305,686 particles were picked in a reference-free manner using Blob Picker, and further refined with Inspect Picks leaving 585,677 particles. 540,665 particles were extracted with a box size of 222 pixels. An initial round of reference-free two-dimensional (2D) classification was performed using 150 classes. The best 13 classes containing 71,628 particles were further classified with 150 classes. The best 12 classes containing 11,519 particles were low-pass filtered to 20 Å and used as templates for a second round of particle picking using Template Picker. 1,579,983 particles were picked and refined to 1,579,983 particles with Inspect Picks. 1,407,670 particles were extracted with an extraction box size of 256 pixels. 2D Classification was used with 200 classes, a maximum resolution of 4 Å, number of online-EM iterations set to 75, with number of final full iterations set to 25, batch size per class of 300, and hard classify for last iteration set to true. The best 16 classes containing 190,512 particles were used to generate an Ab-Initio in C1 using 2 classes. The best class used for Non Uniform Refinement, which generated a 3.17 Å volume in C3. The map was then used for a Reference Based Motion Correction job. Finally, a Non Uniform Refinement was used to generate a: 3.05 Å map in C3 and a 3.35 Å map in C1. Final maps were sharpened with DeepEMHancer^82^. Local resolution estimates were made in CryoSPARC using a fourier shell correlation of 0.143. 3D maps for the two half-maps, the final unsharpened map and the final sharpened map were deposited in the Electron Microscopy Data Bank EMD-48271.

### CryoEM Structure Building and Validation

The PDB model 5JQ3 was used as an initial reference for building the final cryoEM structures. UCSF Chimera^83^ was used to fit the model into density. Multiple rounds of relaxation and minimization were performed on the complete structure, which was manually inspected for errors each time using Isolde^84^, and Coot^85,86^, and Phenix^87^ real-space refinement. Final model quality was analysed using MolProbity^88^. Figures were generated using UCSF ChimeraX^89^. The final structure was deposited in the PDB: 9DZE. For the superposition to 5JQ3, the RMSD calculations were observed in pyMOL using the Align command, with 3945/5825 atoms used for final alignment after 6 cycles.

### Negative Stain Transmission Electron Microscopy

Assembled icosahedral I53-50 GP immunogen samples were thawed and diluted (EBOV-GP-I53-50 sample to 0.05 mg/mL, SUDV-GP-I53-50, EBOV/SUDV-GP-I53-50 – Mosaic, and EBOV-GP-I53-50 + SUDV-GP-I53-50 – cocktail samples to 0.10 mg/mL) in 25 mM Tris/HCl pH 7.5 150 mM NaCl buffer and immediately applied to a freshly glow-discharged 10 nm thick carbon film 400 mesh copper grid (Electron Microscopy Sciences). A volume of 3 µL solution was applied to the grid for 30 seconds before excess liquid was blotted away (Wattman), and immediately replaced with 3 µL 2% Uranyl Formate stain which was subsequently blotted after 10 seconds. Two additional applications and blotting of 2% Uranyl Formate stain were performed preceding the sample being air dried for 5 minutes.

Electron microscopy data was collected for each sample using EPU (ThermoFisher) on a ThermoFisher Talos L120C 120 kV transmission electron microscope with a LaB6 filament and a 2.49 Å pixel size with a CETA camera at 57,000 times magnification. Micrographs were imported into CryoSPARC v4.4.1 (Structura) and Patch CTF Estimation was performed with a minimum resolution of 40 Å and a maximum resolution of 8 Å. Micrographs were blob picked with a minimum diameter of 250 Å and a maximum diameter of 350 Å. Particles were then inspected and those that were accepted were extracted at a box size of 280 pixels (697.2 Å) and binned into 50 2D classes with a batch size of 400 and CTF correction turned off. The best classes for each sample were selected to serve as templates for template picking (with an estimated particle diameter of 280 Å), an additional round of particle inspection, extraction, and an additional round of 2D binning. All jobs in the second round aside from the picking job used the same parameters as they did during the first round. The best classes for each sample were selected for 2D analysis and 3D reconstruction.

### Mouse Immunizations and maEBOV Viral Challenge

At 10-15 weeks of age, 10 female BALB/c mice (Jackson Laboratories) per dosing group were vaccinated by the subcutaneous (SC) route with a prime immunization, and three weeks later mice were boosted with a second vaccination SC. Prior to inoculation, immunogen suspensions were gently mixed 1:1 vol/vol with AddaVax adjuvant (Invivogen, San Diego, CA) to reach a final concentration of 0.01 mg/mL antigen. To obtain sera all mice were bled two weeks after prime and boost immunizations. Blood was collected by the submandibular or retro-orbital route and rested in 0.8 mL serum separator tube (SST) tube at room temperature for 10 min to allow for coagulation. Serum was separated from red blood cells via centrifugation at 10,000 g for 10 min. Serum was stored at 4°C or 80°C until use. For viral challenge, mice were acclimated to ABSL-4 for at least 7 days and were injected by the intraperitoneal (IP) route with a 100 pfu target dose of maEBOV^57^ in a volume of 200 µL. Post-exposure, mice were observed daily for weight and clinical signs of disease for up to 28 days. Upon onset of clinical signs of disease, mice were observed at least twice daily. Moribund mice, with maximum clinical disease scores of 4 (defined as bleeding, unresponsive, weak, unable to walk) or surviving through day 28 post exposure were euthanized.

### Guinea Pig Immunizations and gpSUDV Viral Challenge

Groups of 6 female Hartley guinea pigs (Charles River) per dosing group (except for Group 1 which had 5 animals due to one loss in preparation), weighing 350-500 g were vaccinated with a prime immunization, and three weeks later guinea pigs were boosted with a second vaccination by the IP route. Prior to inoculation, immunogen suspensions were gently mixed 1:1 vol/vol with AddaVax adjuvant (Invitrogen, San Diego, CA) to reach a final concentration of 0.01 mg/mL antigen. To obtain sera all guinea pigs were bled two weeks after prime and boost immunizations. Blood was collected via submental venous puncture and rested in 0.8 mL SST tube at room temperature for 10 minutes to allow for coagulation. Serum was separated from red blood cells via centrifugation at 10,000 g for 10 minutes. Complement factors and pathogens in isolated serum were heat-inactivated via incubation at 56°C for 60 minutes. Serum was stored at 4°C or 80°C until use. At approximately 8 weeks post-prime vaccination, animals were moved into ABSL-4 for acclimation. At ABSL-4, guinea pigs were exposed by the IP route with 1000 pfu of gpaSUDV^61^ in a volume of 400 µL. Guinea pig clinical observations, weights, and temperatures (by subcutaneous chip; BDMS) were recorded daily for 28 days. Moribund guinea pigs were promptly euthanized when they met predefined euthanasia criteria, a score of 4; defined as bleeding, unresponsive, weak, or unable to walk.

### ELISA

Endpoint IgG serum ELISAs to detect EBOV or SUDV antibodies were conducted using the following assay. ELISA plates (Falcon) were coated with recombinant SUDV-GP or recombinant EBOV-GP (NCI, Frederick MD) at 0.1 µg/well in phosphate buffer saline (PBS) (Sigma-Aldrich) and stored at 4°C overnight. After blocking with PBS/5% Milk (BioWorld) /0.05% Tween 20 (Sigma) (blocking buffer) at room temperature (RT) for 2 hours, pre-diluted serum samples and positive and negative control samples, all diluted in blocking buffer, were added to wells in duplicate. Following 1 hour incubation at RT, plates were washed with PBS/0.05% Tween 20. Secondary antibody (goat anti-mouse or guinea pig IgG-HRP) (KPL) diluted in blocking buffer was added to the plates at RT for 1 hour. After washing plates with PBS/0.05% Tween 20, 2,2’-azino-bis(3-ethylbenzothiazoline-6-sulfonic acid (ABTS) (SeraCare) was added for 30 minutes at RT and absorbance was measured at 405 nm using a SpectraMax® M5 (Molecular Probes). The reciprocal endpoint dilutions were reported.

### Statistics and Reproducibility

Statistical details of experiments can be found in the figure legends. For animal samples analysis, no blinding of the experimenter was done. For animal studies, 10 BALB/c or 6 guinea pigs were used. No statistical methods were used to predetermine sample size. For animal samples analysis, no blinding of the experimenter was done. Geometric mean anti-GP IgG titers were calculated based on absolute ELISA signals with background removed. For the ELISA plot data in the maEBOV study we ran statistical tests and calculated significance levels by comparison of all individual test article Groups to the bare I53-50 nanoparticle group. We first conduct normality testing using the Shapiro-Wilk test to assess normality for both the current Group and the reference Group 8. If both groups are normally distributed (p > alpha), we perform an independent t-test. If not, we use the Mann-Whitney U test as a non-parametric alternative. We assume normality since Shapiro-Wilk requires at least 3 observations, and all groups had more than 3 data points. We determined the significance level based on the P value. The level of statistical significance is indicated as: *P < 0.05; **P < 0.01; ***P < 0.001. Kruskal–Wallis survival tests were performed for survival data, and calculated P values for statistical significance are indicated for comparisons against the respective naked I53-50 NP control group based on log-rank Mantel-Cox statistical analyses. The level of statistical significance is indicated as: *P < 0.05; **P < 0.01; ***P < 0.001.

Production of EBOV and SUDV GP constructs reported was produced at least once using lentivirus production and at least twice using pCMV/R transient transfection. IMAC and SEC purification and analytical HPLC SEC was performed for all constructs and nanoparticle preparations at least once for in vitro characterization and multiple times prior to animal studies. Representative SEC profiles and SDS-PAGE analyses for various constructs were selected for comparison. Production and certificate of testing for all GP-I53-50 nanoparticle immunogens was completed on at least two separate occasions for the mouse and guinea pig studies. Negative-stain EM was performed routinely for all GP-I53-50 nanoparticle assemblies during in vitro characterization and production of immunogens for animal studies. All ELISA binding assays were performed in duplicate. Due to the limited availability of purified mouse or guinea pig IgG samples, in vitro viral neutralization assays were not performed.

## Acknowledgements

The project or effort depicted was or is sponsored by the Defense Threat Reduction Agency Grant HDTRA1-18-1-0001 (D.B., N.P.K, L.J.S., S.P.W., J.M.D., R.R.B., S.E.Z., and E.E.Z.), the Audacious Project at the Institute for Protein Design (L.C., M.M., R.R., S.C., C.E.C., C.W., A.J.B., R.S., B.F., and K.D.C.) and The Open Philanthropy Project for Universal Flu Vaccine and Improving Protein Design (N.B., N.P.K., and D.B.). The authors wish to thank Jane Carter for her support in production and purification of lentivirus produced immunogens. Figure 3a was created with biorender.com and ChimeraX.

## Author Contributions

L.J.S., D.B., J.M.D., A.J.B. and N.P.K. designed and supervised the research. C.E.C., M.M., R.R., and S.C., engineered expression vectors, produced and purified various glycoprotein immunogen constructs. N.B., S.P.W., B.F., L.C., produced, purified, and characterized the biophysical properties of all I53-50 nanoparticle immunogens. C.W., A.J.B., R.S., B.F., and K.D.C., prepared samples for negative stain and cryo electron microscopy and structurally characterized the various antigens, nanoparticle components, and assembled nanoparticle immunogens detailed in this study. S.E.Z., E.E.Z., and R.R.B. performed all animal studies in BSL-4 and serological analyses of IgG responses to nanoparticle immunogens. L.C. and L.J.S. coordinated the resources and funding required for the research. L.J.S., C.W., and N.B. drafted and wrote the manuscript. L.J.S., C.E.C., C.W., N.B., S.E.Z., and E.E.Z. prepared figures for the manuscript. All authors analyzed data and revised the manuscript.

## Data Availability

The datasets generated during and/or analyzed during the current study are available from the corresponding author upon reasonable request. The atomic coordinates and map for our CryoEM structure of the perfusion EBOV-GP-Foldon construct is located at the Protein Data Bank PDB ID: 9MHA, EMDB ID: EMD-48271.

## Disclaimers

The opinions, interpretations, conclusions, and recommendations presented are those of the authors and are not necessarily endorsed by the U.S. Army or Department of Defense. The funders had no role in the design of the study; in the collection, analyses, or interpretation of data; in the writing of the manuscript; or in the decision to publish the results. The use of trade or manufacturers’ names in this publication does not constitute an official endorsement of any commercial products. This report may not be cited for purposes of advertisement. Contractor affiliation of some authors does not imply endorsement by the U.S. Government of any contractor.

## Institutional Review Board Statement

All animal studies were conducted under IACUC-approved protocols in compliance with the Animal Welfare Act, Public health Service Policy, and other applicable federal statutes and regulations relating to animals and experiments involving animals. The facility where these studies was conducted (USAMRIID) is accredited by the Association for Assessment and Accreditation of Laboratory Animal Care, International and adheres to principles stated in the Guide for the Care and Use of Laboratory Animals, National Research Council, 2011. The animal study protocol was approved by the Institutional Animal Care and Use Committee of USAMRIID (protocol number AP-19-005, 05 February 2019).

